# A quantitative description of light-limited cyanobacterial growth using flux balance analysis

**DOI:** 10.1101/2024.01.10.574812

**Authors:** R. Höper, D. Komkova, T. Zavřel, R. Steuer

**Affiliations:** Institute for Biology, Theoretical Biology (ITB), Humboldt-University of Berlin, Philippstr. 13, Haus 20, 10115 Berlin,Germany; Global Change Research Institute of the Czech Academy of Sciences, Bělidla 986/4a, Brno, 603 00,Czechia

**Author notes:** Contributing authors.

**Keywords:** *Synechocystis* sp. PCC 6803, metabolic reconstruction, constraint-based analysis, FBA, bioenergetics, photosynthesis, photobioreactor, quantum yield, systems biology

## Abstract

The metabolism of phototrophic cyanobacterial is an integral part of global biogeochemical cycles, and the capability of cyanobacteria to assimilate atmospheric CO_***2***_ into organic carbon has manifold potential applications for a sustainable biotechnology. To elucidate the properties of cyanobacterial metabolism and growth, computational reconstructions of the genome-scale metabolic networks play an increasingly important role. Here, we present an updated reconstruction of the metabolic network of the cyanobacterium *Synechocystis* sp. PCC 6803 and its analysis using flux balance analysis (FBA). To overcome limitations of conventional FBA, and to allow for the integration of quantitative experimental analyses, we develop a novel approach to describe light absorption and light utilization. Our approach incorporates photoinhibition and a variable quantum yield into the constraint-based description of light-limited phototrophic growth. We show that the resulting model is capable to predict quantitative properties of cyanobacterial growth, including photosynthetic oxygen evolution and the ATP/NADPH ratio required for growth and cellular maintenance. Our approach retains the computational and conceptual simplicity of FBA and is readily applicable to other phototropic microorganisms.

## 1 Introduction

Oxygenic photosynthesis is one of the most important biological processes on our planet and drives primary production in almost all ecosystems. To this day, cyanobacteria, the evolutionary inventors of oxygenic photosynthesis, remain an integral part of global biogeochemical cycles. In addition, because of their capability to assimilate atmospheric CO_2_ into organic carbon using sunlight as the only source of energy, cyanobacteria are an interesting resource for green biotechnology. Among cyanobacteria, the strain *Synechosystis* sp. PCC 6803 is an established model organism with a broad compendium of published studies that characterize its growth and metabolism under different environmental conditions [1, 2].

Concomitant to experimental studies, computational reconstructions of cellular metabolism play an increasing role to allow us to understand cyanobacterial physiology and to predict properties of light-limited metabolism and growth. Genome-scale metabolic reconstructions (GSMRs) are available for an increasing number of microbial organisms, including *Synechosystis* sp. PCC 6803 [3–8] and several other cyanobacteria [9–13].

GSMRs aim to provide a comprehensive account of the stoichiometry of the metabolic reaction network within a microbial organism. The construction of GSMRs is typically based on an available genome sequence. Annotated genes and the encoded protein complexes are linked to suitable reaction databases, such as KEGG [14, 15] or MetaCyc [16] to establish gene-protein-reaction relationships. A GSMR includes all known enzyme-catalyzed metabolic reactions, transport reactions, as well as non-catalyzed processes, such as diffusion or spontaneous degradation of metabolites.

Once established, a number of computational techniques are available to analyze a GSMR and investigate its metabolic capabilities under different environmental conditions. In particular, methods based on linear programming (LP), such as flux balance analysis (FBA) [17, 18], have become a de-facto standard to analyze GSMRs. The success of FBA is due to its computational simplicity, as well the fact that the application of FBA only requires knowledge of the stoichiometry of the metabolic network, a suitable objective function, and a set of constraints on uptake fluxes–and does not require extensive knowledge about enzyme-kinetic parameters or regulatory interactions. Instead, predictions using FBA are based on the assumption of evolutionary optimality. That is, FBA seeks to predict maximal growth rates and the associated metabolic fluxes based on the assumption that an organism maximizes its growth rate given the constraints on nutrient uptake rates set by the environment [19, 20]. While the conditions under which the assumption of evolutionary optimally holds is still subject to considerable debate, predictions are often in good agreement with available data [20–22].

Different from heterotrophic metabolism, however, the description of photoautotrophic metabolism gives rise to additional challenges due to the use of light as the primary source of energy. Light absorption, photodamage, and the unique redox metabolism associated with photosynthesis are key aspects in a computational description of photoautotrophic growth. Compared to heterotrophic growth, only few studies provide a quantitative computational analysis of light-limited phototrophic growth and integrate physiologically relevant photosynthetic properties into large-scale models of light driven metabolism [23].

The purpose of this work is to provide an updated GSMR of the model cyanobacterium *Synechosystis* sp. PCC 6803 and its quantitative analysis using previously published growth experiments [1]. Specifically, we seek to describe light-limited growth of the cyanobacterium *Synechocystis* sp. PCC 6803 based on a genome-scale metabolic reconstructions that incorporates key photosynthetic properties and energy flows. To this end, we introduce a novel approach to describe light absorption in FBA and to account for the effects of photoinhibition. We show that the resulting description is capable to predict quantitative properties of phototrophic growth, in particular photosynthetic oxygen release, as well as the ratio between linear and cyclic electron transport. The results demonstrate that current GSMRs, together with appropriate constraints, are suitable to describe and predict quantitative aspects of phototrophic growth. Our method is based on only few additional parameters that have a clear interpretation in the context of phototrophic growth, and whose numerical values can be determined from a measured growth-irradiance curve. We further provide a stoichiometrically reduced core model for applications that benefit from a reaction network that contains fewer reactions but nonetheless remains stoichiometrically correct.

## 2 Results and Discussion

### 2.1 Network reconstruction and FBA

The metabolic model of the cyanobacterium *Synechocystis* sp. PCC 6803 is based on previously published reconstructions, in particular Knoop et al. (2013) and Knoop et al. (2015) [5, 24]. The current reconstruction includes revised gene-protein-reaction associations, revised stoichiometric balances, and an increased coverage of metabolic processes. Additional reactions include the Entner–Doudoroff pathway [25], enabling the oxidation of glucose to gluconate and its subsequent phosphorylation to 6-phosphogluconate, as well as additional reactions for biotin metabolism and purine and pyrimidine metabolism. Additions are based on primary literature, as well as on a comparison with the KEGG database (reference genome BA000022) [26]. A detailed list of added reactions and their associated genes, compared to Knoop et al. (2015) [24], is provided as Supplementary File S5.

The reconstruction consists of a total of 831 reactions, 715 metabolites (615 unique metabolites), and covers 796 genes. The 831 reactions include 778 mass balanced intracellular reactions, as well as the supply of nutrients into the growth medium (denoted as intracellular space in Figure 1) and biochemical interconversion therein, such as CO_2_ to 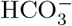. Metabolites and reactions are organized into 7 compartments: extracellular space (growth medium), periplasm, cytoplasmic membrane, cytoplasm, carboxysome, thylakoid membrane, and thylakoid lumen, see Figure 1. The model was tested for stoichiometric consistency using MEMOTE [27]. Details of the network reconstruction and analysis are provided in the Materials and Methods. An annotated SBML file is available as Supplementary File S2. In addition we provide a reduced model (Supplementary File S4) using an algorithm for stoichiometric network reduction [28], see Materials and Methods.

**Fig. 1:**
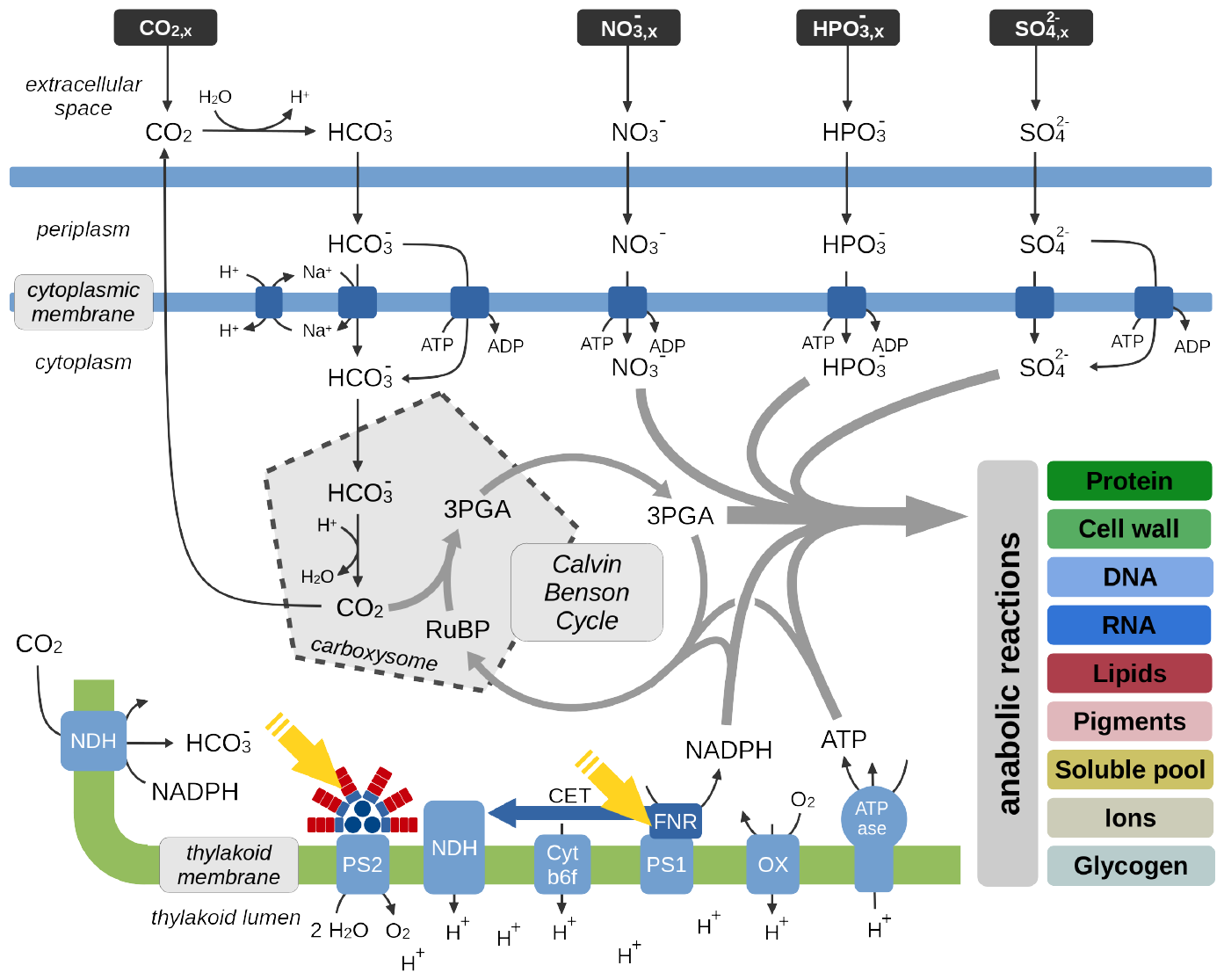
Overview of key metabolic processes involved in cyanobacterial phototrophic growth and described in the metabolic reconstruction of *Synechocystis* sp. PCC 6803. Growth of *Synechocystis* sp. PCC 6803 is characterized by the net uptake of bicarbonate 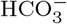 from the extracellular space. Within the carboxysomes, 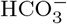 is converted into CO_2_ by a carbonic anhydrase and assimilated by the ribulose-1,5-bisphosphat-carboxylase/-oxygenase (RuBisCO). Organic carbon is utilized to synthesize cellular biomass using *ATP* and *NADPH* re-generated by the photosynthetic light reactions. Cellular biomass consists of protein, cell wall components, DNA, RNA, lipids, pigments, a soluble pool (metabolites and cofactors), ions, and glycogen as a storage component. The reconstruction contains 778 mass-balanced intracellular reactions and covers 796 genes. Metabolites are organized into 7 compartments. Respiratory complexes in the cytoplasmic membrane are not shown. Abbreviations: ribulose-1,5-bisphosphate (RuBP), glyceraldehyde-3-phosphate (3PGA).

Analysis of a GSMRs using FBA typically requires the definition of a biomass objective function (BOF). The BOF specifies the amounts of cellular components needed for the synthesis of one gram cellular dry mass (gCDM). In the following, we use the (static) BOF defined in previous reconstructions as a reference [3, 5], and compare the results with an experimentally obtained light-dependent BOF. The latter is derived from measurements of the mass fractions of protein and glycogen as a function of light intensity, with the remaining components scaled accordingly. Figure 2 summarizes the available growth data [1], including the specific growth rate as a function of light intensity, the respective changes in biomass composition, as well as the light-dependent oxygen (O_2_) evolution. All data are sourced from a single set of experiments [1].

**Fig. 2:**
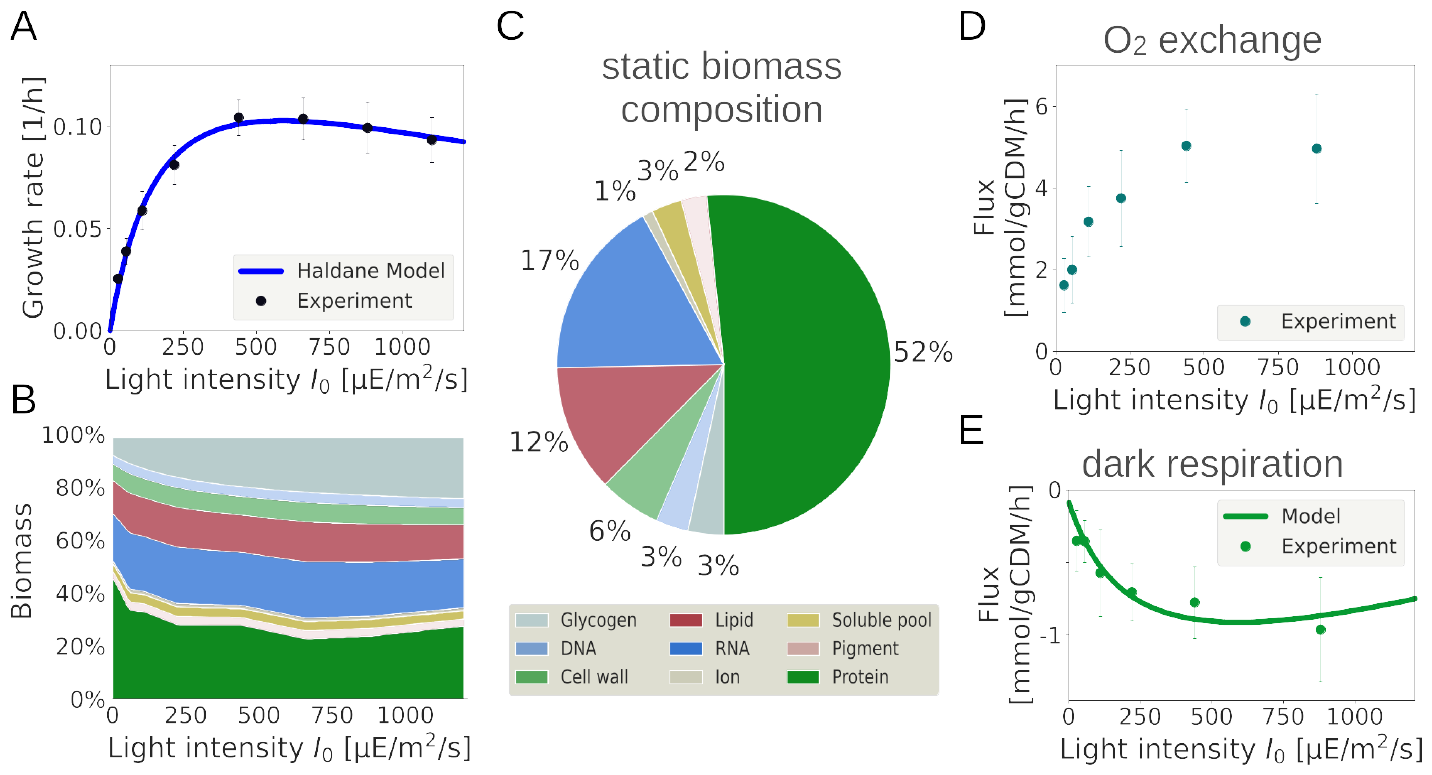
Physiological properties of phototrophic growth of *Synechocystis* sp. PCC 6803 obtained in a light-limited turbidostat [1]. (A) The specific growth rate as a function of light intensity. The functional form of the growth rate is consistent with a phenomenological Haldane/Aiba equation (solid line). (B) The light-dependent biomass composition (BOF), derived from the measured protein and glycogen mass fractions. (C) A static biomass composition as a reference, as used in previous analyses. (D) Measured net oxygen (O_2_) evolution as a function of light intensity. (E) Measured oxygen (O_2_) consumption shortly after stopping illumination as a function of light intensity. The measured O_2_ consumption in darkness is used as a proxy for respiration during illumination, and is part of parameter estimation. The solid green line shows the O_2_ consumption of the fitted model. Growth data were originally described in Zavřel et al. [1] and serve here to parameterize and test the model. In the following, we use *µE* as an abbreviation for *µmol photons* in the description of light intensity.

Using the light-dependent BOF, the metabolic reconstruction of *Synechocystis* sp. PCC 6803 allows for phototrophic growth with atmospheric CO_2_ as sole source of carbon (taken up as bicarbonate 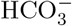 from the extracellular medium). When maximizing the BOF with nitrate 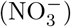 as the sole source of nitrogen, the metabolic reconstruction gives rise to the following overall stoichiometry for the synthesis of one gram cellular dry mass,

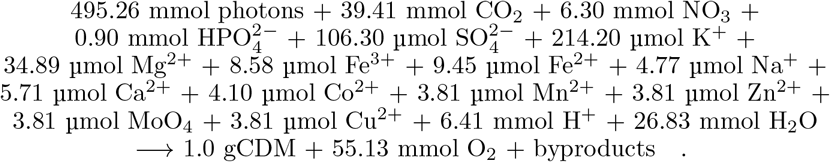

The overall stoichiometry is estimated using the BOF for a light intensity *I*_0_ = 660 *µ*mol (photons) m^*−*2^ corresponding to a growth rate of 0.10 h^*−*1^ with a mass fraction of 23% protein and 20% glycogen. The overall stoichiometry indicates that (maximal) biomass yield necessitates the excretion of three carbon-containing byproducts. The existence of such obligatory byproducts points to either errors or inaccuracies in the current reconstruction, or the existence of biologically inevitable side products that are not recycled within metabolism (see Supplementary Table S2 for a list of products). The byproducts, however, constitute less than 0.07‰ of gCDM, and are therefore unlikely to be linked to exudation of compounds observed for cyanobacterial cells [29].

According to the overall stoichiometry, carbon constitutes approximately 47% of cellular (dry) mass, the C:N content is approximately 6.3:1 (slightly lower than the Redfield ratio of approximately 6.6:1, but slightly larger than the experimentally determined ratio, see Supplementary Figure S5). The photosynthetic quotient *PQ*, the ratio of *O*_2_ production relative to CO_2_ assimilation is PQ ≈ 1.4, in good agreement with experimental analysis [1, 5]. Considering only the stoichiometric requirements for the synthesis of biomass, that is, neglecting cellular maintenance and other processes not related to formation of biomass, such as photorespiration, the maximal yield of biomass per photon is 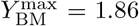 gCDM/mol photons for the static reference BOF, and and ranges from 1.89 to 2.03 gCDM/mol photons for the light-dependent BOF, similar to values reported for metabolic reconstructions of other cyanobacteria [30, 31]. The differences are primarily due to the varying mass fraction of protein and glycogen in the light-dependent BOF. Stoichiometrically, minimally ≈ 9 photons are required to produce one molecule *O*_2_ during growth, slightly lower than the experimentally determined values of 11 photons per *O*_2_ produced [32].

### 2.2 Modeling light absorption and utilization in FBA

Our aim is a quantitative description of light-limited phototrophic growth. In particular, we seek to reproduce the measured specific growth rates of the cyanobacterium *Synechocystis* sp. PCC 6803 over a wide range of light intensities, as obtained by an experimental evaluation using a highly controlled and reproducible cultivation setup [1].

As shown in Figure 2, and characteristic for phototrophic microorganisms [31, 33, 34], the light-limited growth rate as a function of the light intensity *I* (measured in mol photons per area per time) can be described by a Haldane or Aiba equation,

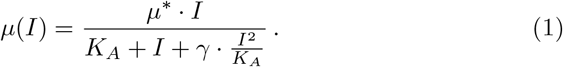

The phenomenological Haldane/Aiba equation is specified by three parameters, *µ*^*∗*^,*K*_*A*_, and *γ*.

The latter is a dimensionless parameter that quantifies the inhibitory impact of light on the growth rate (photoinhibition).

In contrast, within constraint-based models of phototrophic growth, the uptake and utilization of photons is typically described analogous to the uptake of nutrient molecules and, in the absence of further constraints, the growth rate scales linearly with the rate of photon uptake *J*_*I*_ (measured in mol photons per gCDM per time). In the following, we therefore propose a novel approach to incorporate light uptake and utilization into constraint-based models of phototrophic growth–with the aim to better capture the growth properties of cyanobacteria and other phototrophic microorganisms. Our approach is motivated by mechanistic models of phototrophic growth [31]. Specifically, we consider a two step process in which photons are first absorbed by the cell and are then utilized with a light-dependent *photosynthetic efficiency* (also denoted as light-dependent *quantum yield*).

Following the experimental setup of Zavřel et al. [1], we first consider a light source with a single wavelength (monochromatic light). The rate of photon flux is described by an incident light intensity or photon flux density *I*_0_ (mol photons per m^2^ per second) at the surface of the culture vessel. Photons are absorbed by the culture with a proportionality factor *α* (measured in m^2^ per gCDM). The factor *α* describes an effective area of light absorption per gCDM. The rate *J*_*I*_ of photon uptake or photon absorption is

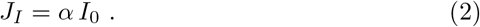

Photons are then utilized with a (dimensionless) quantum yield *η*(*J*_*I*_). The quantum yield describes the ratio of the rate of utilized photons 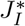 relative to the rate of absorbed photons *J*_*I*_,

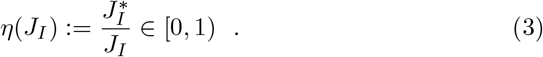

Following mechanistic models of light utilization [31], we postulate that the quantum yield is a decreasing function of the rate of photon uptake,

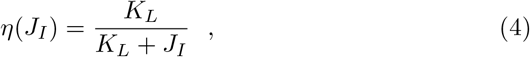

with *K*_*L*_ as an adjustable parameter. A formal derivation of equation (4) based on a 2-state model of photosynthesis is provided in the Materials and Methods. Analysis of the constraint-based model is then based on an effective rate of light uptake 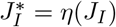. *J*_*I*_ (measured in mol photons per gCDM and time),

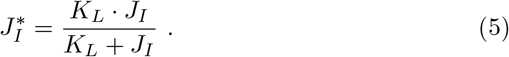

The effective rate of light uptake 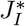 serves as a constraint for the maximization of the BOF in FBA, and depends on the parameters *α* and *K*_*L*_, as well as on the incident light intensity *I*_0_. The parameter *K*_*L*_ can be interpreted as a maximal capacity of light utilization, i.e., the maximal number of photons the cell can utilize per gCDM and time. Specifically, equation (4) implies that under low light conditions, almost all absorbed photons are utilized, 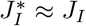 for *J*_*I*_ ≪ *K*_*L*_, whereas for high light intensities there is an upper limit to the rate of light utilization, 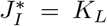 for *J*_*I*_ → ∞. Hence, together with the maximal biomass yield 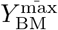, the parameter *K*_*L*_ also provides an upper bound for the maximal growth rate in the absence of photoinhibition or other detrimental factors.

Both parameters, *α* and *K*_*L*_, may depend on the specific strain and culture conditions and can be estimated from the experimental growth-irradiance curve shown in Figure 2A.

### 2.3 Describing photoinhibition

To account for photoinhibition, we include a description of light-induced photodamage, in particular of the D1 protein of photosystem II. Since, within our approach, protein turnover is not explicitly modelled, we describe photoinhibition by a light-dependent ATP utilization that accounts for the increased repair and translation mechanisms in dependence of light. We emphasize that photodamage is a well studied phenomenon, and there is broad experimental evidence across multiple domains of life, from cyanobacteria to eukaryotic algae and plants, that photodamage occurs at all light intensities (i.e., not only at high light intensities) and is proportional to the light intensity [35–39]. Hence, to account for photodamage, we introduce an light-dependent rate *ν*_*D*_ of ATP utilization as an additional constraint in our analysis,

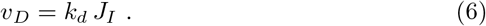

The rate *ν*_*D*_ is proportional to the rate *J*_*I*_ of light absorption, with a (dimensionless) proportionality factor *k*_*d*_, and accounts for the increased ATP demand as a consequence of increased protein turnover and repair.

Within the model, the description of light absorption, utilization, and photodamage therefore has three adjustable parameters, *α, K*_*L*_, and *k*_*d*_, analogous to the three parameters of the phenomenological Haldane equation (1). Below, we demonstrate that these parameters can be determined from a fit of the constraint-based model to the measured specific growth rate over the full range of light intensities. Once the numerical values of the parameters are known, additional growth properties can be evaluated.

### 2.4 Incorporating blue background illumination

Prior to a numerical analysis, however, we have to account for the specific experimental setup used in Zavřel et al. [1]. In addition to monochromatic red light (*λ*_max_ ≈ 633 nm), supplied with an intensity *I*_0_ between 27.5 and 1100 *µ*mol (photons) m^*−*2^ s^*−*1^, the growth setup was supplemented with blue light (*λ*_max_ ≈ 445 nm) with a constant intensity *I*_*b*_ = 27.5 *µ*mol (photons) m^*−*2^ s^*−*1^. The reason for the additional background illumination was to avoid possible (regulatory) growth limitations resulting from the absence of short wavelength photons [1].

In the following, we therefore make use of a heuristic *ansatz* that accounts for the utilization of blue light, while avoiding additional complexity in the model description. Instead of equation (2), the rate of photon absorption is described by

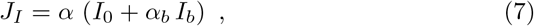

where *α*_*b*_ quantifies the relative impact of the additional (and constant) blue background intensity. Equation (7) postulates that the blue background light contributes to the overall light intensity with a weight *α*_*b*_. Computational details are provided in the Materials and Methods.

### 2.5 Additional physiological flux constraints

When estimating the maximal growth rate and other physiological properties, constraint-based models are typically subject to additional constraints that ensure biologically plausible solutions. That is, these additional constraints are part of the model definition and can not be derived from the simulation experiments.

Following previous studies [5, 24], we assume that growth of *Synechcystis* sp. PCC 6803 is subject to a set of additional flux constraints summarized in Table 1: firstly, we enforce a (small) flux through the ribulose-1,5-bisphosphate-oxygenase, i.e., the use of molecular oxygen O_2_ by RuBisCO instead of CO_2_ as a substrate, resulting in no net CO_2_ fixation and the formation of phosphoglycolate as a by product (photorespiration). The flux is set to 3% of the total RuBisCO flux. Secondly, we enforce a non-zero flux through the Mehler and Mehler-like reactions (the former play a minor role in cyanobacteria). The values are listed in Table 1 and are chosen according to values used in previous recontructions [5]. Thirdly, we enforce a non-zero flux through the terminal oxidase and a non-growth associated maintenance (NGAM) reaction, accounting for basal ATP utilization that is not associated with the synthesis of biomass.

**Table 1:**
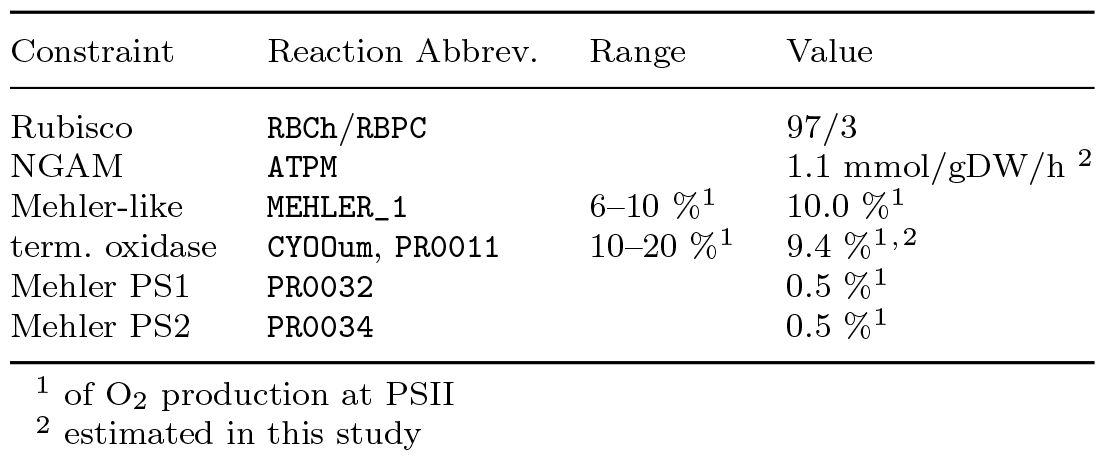
Additional physiological constraints used in the simulations to ensure biologically plausible solutions. Values are adopted from previous studies [5], except otherwise noted. All percentages are relative to O_2_ production at PSII.

In previous analysis [5], the (lower bound of) flux through the terminal oxidase was assumed to be 10% of O_2_ production at PSII, the NGAM reaction was assumed be 1.3 mmol/gDW/h, approximately 10% of maximal ATP production. In the following, however, we consider both parameters as unknown and adjust the values using the data reported by Zavřel et al. [1].

Specifically, in addition to the O_2_ evolution as a function of light intensity, the O_2_ consumption shortly after onset of darkness was measured (Figure 2D). The measured O_2_ consumption in darkness varies as a function of the previous light intensity and can serve as a proxy for O_2_ consuming processes that are not directly related to PSII activity. That is, we assume that the observed non-light associated O_2_ consumption is also present during illumination. Within the model, non-light associated O_2_ consumption is defined as flux through the terminal oxidase, as well as O_2_ that is used as a substrate in metabolic reactions. Hence, non-light associated O_2_ consumption can be calculated as O_2_ evolution at PSII minus net O_2_ export into the extracellular medium minus flux through the Mehler-like reactions (the latter are assumed to stop rapidly when illumination stops). We note that the exact values of the flux through terminal oxidase and NGAM have no major impact on the results reported below.

Finally, based on the experimental characterization of the cellular constituents by Zavřel et al. [1], in particular the light-dependent changes in glycogen and protein content, we adjust the biomass composition for each light intensity. The results are compared to solutions obtained for the constant reference BOF (see Figure 2B and Supplemental Figure S4).

### 2.6 Estimation of growth parameters

Given the physiological constraints on the flux distribution listed in Table 1, we can now estimate the 4 parameters used in the description of light absorption and utilization, *α, α*_*b*_, *K*_*L*_, and *k*_*d*_, as well as the values of 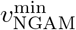 and 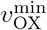, such that the maximal specific growth rates matches the experimental values shown in Figure 2A, and the non-light associated O_2_ consumption matches the O_2_ consumption after illumination stops (Figure 2D). Computational details are described in the Materials and Methods.

Figure 3 shows the growth rate as a function of light intensity when the description of light absorption and utilization is iteratively refined. Starting with equation (7) only, and using *α* and *α*_*b*_ as adjustable parameters, the growth rate is a linear function of (red) light intensity with a slope that matches the growth curve at low light intensities. Including the photosynthetic efficiency *η*(*J*_*I*_), equation (3), with *K*_*L*_ as an additional free parameter, results in a saturation of the growth rate as a function of light intensity. With the addition of photodamage (*k*_*d*_ as an additional parameter), we obtain the final fit of the growth curve across the entire range of light intensities. The estimated parameters are *α* = 0.13 ± 0.01 [m^2^*/*gCDM], *α*_*b*_ = 0.63 ± 0.24 [n.d.], *K*_*L*_ = 115.00 ± 12.8 [mmol(photons)*/*gCDM*/*h], and *k*_*d*_ = 0.07 ± 0.018 [n.d.].

**Fig. 3:**
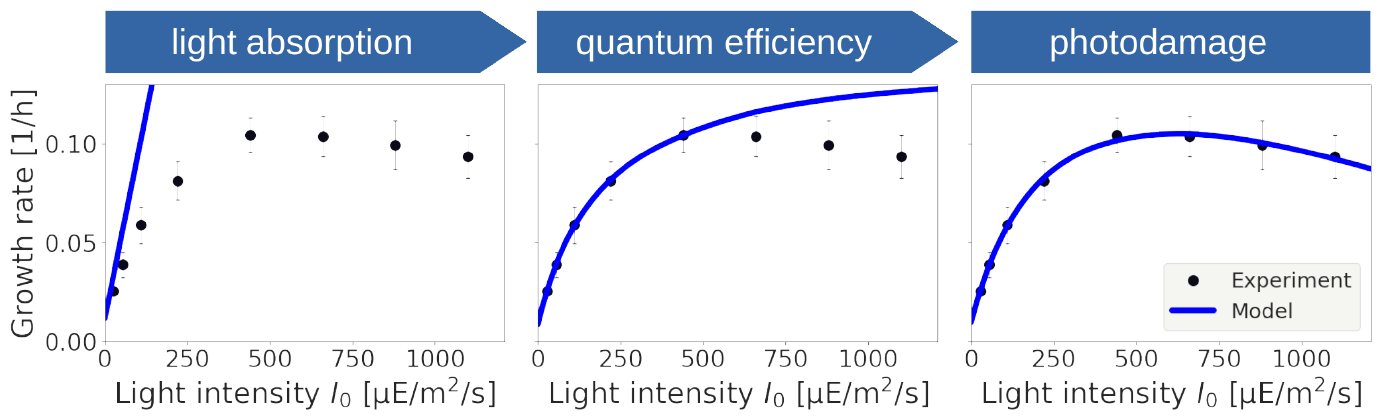
Iterative refinement of the description of the specific growth rate as a function of the light intensity. Incorporating only light absorption (equation (7)) results in a linear dependence of the growth rate on the light intensity, analogous to conventional FBA. Incorporating a variable quantum yield (equation (3)) results in saturating growth with a maximal photosynthetic capacity *K*_*L*_. Finally, including photoinhibition (equation (6)) results in the final quantitative fit of the model to experimental values.

The estimated parameters for the terminal oxidase and the NGAM reaction are 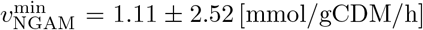 and 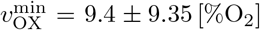. The estimated values are very close to values used in previous reconstructions [5], and are subject to substantial error intervals. The latter is expected, as ATP utilization for maintenance (NGAM) is known to be small in *Synechocystis* 6803 and the value is primarily estimated from the (offset of the) growth rate at low light intensities.

We note that the detailed composition of the BOF has no major impact on the quality of the fit (see Supplementary Figure S1 for a fit using the static reference BOF). The latter is consistent with previous studies showing that the detailed composition of the BOF has only a weak impact on the estimated growth rate [40]. We further note that the fitted values are specific for the strain and the experimental setup and are not necessarily universal parameters of phototrophic growth. Nonetheless, we can interpret the numerical values in the context of our analysis.

Firstly, as discussed above, the parameter *K*_*L*_ can be interpreted as a maximal capacity of light utilization. Together with the (maximal) biomass yield *Y* ≈ 1.9 to 2.0 gCDM/mol (photons) the capacity suggests a maximal growth rate of up to *µ*^max^ = 0.23 h^*−*1^, corresponding to a minimal division time of approximately 3.0 h in the absence of photoinhibition and other detrimental effects. The value is slightly lower than the minimal division time reported for *Synechcystis* sp. PCC 6803 observed under optimal conditions [41], but plausible given that photoinhibition and other detrimental effects cannot be fully alleviated.

Secondly, the estimated value of *α*_*b*_ ≈ 0.63 is in agreement with experimental previous studies. Blue light is predominately absorbed at PSI and was shown to less efficient for Synechocystis growth compared to red light [42]. For example, it has been shown that an increase of orange/red photons (633 ± 20 nm) from 110 to 220 *µ*mol (photons) m^*−*2^ s^*−*1^ increased the specific growth rate by 29%, whereas the same addition of blue photons (445 ± 20 nm) increased growth rate by 14% only [43]. Likewise, cultivating Synechocystis under blue light alone resulted in reduction of maximum specific growth rate by 50– 75% [42], compared to growth under orange/red light [44]. An estimated value of *α*_*b*_ ≈ 0.63 reflects the observed lesser contribution of blue light to growth.

### 2.7 Comparison with a mechanistic model of light uptake

The assumption that light uptake is directly proportional to the incident light intensity, as assumed in equations (2) and (7), does not correspond to a mechanistic model of light absorption. Rather, our approach is motivated by the fact that it can be readily applied as long as a measured dependency of growth rate on light intensity is available, even when detailed information about the light spectrum and geometry of the reactor vessel are lacking.

The use of a flat panel photobioreactor in Zavřel et al. [1], however, allows us to compare the results with a mechanistic description of light absorption that takes reactor geometry into account. Specifically, neglecting deviations due to scattering and fluorescence, light attenuation as a function of reactor depth can be approximated by the Lambert-Beer equation. In a flat-panel reactor the light intensity at depth *z* is approximately *I*(*z*) = *I*_0_ exp (− *ϵ ρV z*), where *I*_0_ denotes the incident light intensity at the surface, *ϵ* the absorption coefficient (in units m^2^/gCDM), and *ρV* the volumetric density (in units gCDM/m^3^). Integrating over a flat panel reactor of depth *Z* then results in a total rate 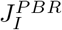 of absorbed photons [30, 31, 45],

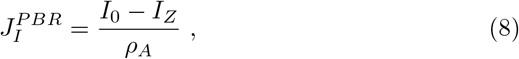

where *I*_*Z*_ is the light intensity at depth *Z* (transmitted light) and *ρA* = *ρV* · *Z* is the areal biomass density (units: gCDM/m^2^).

Figure 4 compares the estimated photon flux 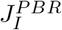 to the photon flux previously obtained from equation (7). In Figure 4 the contribution of the background blue light to 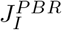 is assumed to range from 0 to 100% and is shown as a (small) shaded area. That is, we assume the absorption coefficient for blue light is identical to red light and the photons are added to the red light intensity with a weight from 0 to 100% (the contribution is only visible at low light intensities). The remaining variability in 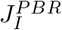 is primarily due to uncertainty in the estimation of the areal biomass density. Both values for the photon flux are in good agreement, implying that equation (7) results in a similar amount of absorbed photons as the estimated flux using reactor geometry. For low light intensities, the absorbed photons slightly exceed the amount fitted in equation (7). For high light intensities, however, equation (7) assumes a slightly higher photon utilization rate as actually absorbed. To resolve the small discrepancy, we note that photodamage is modelled as ATP utilization only, and therefore might overestimate the ATP, and hence photon requirements, at high light intensities.

**Fig. 4:**
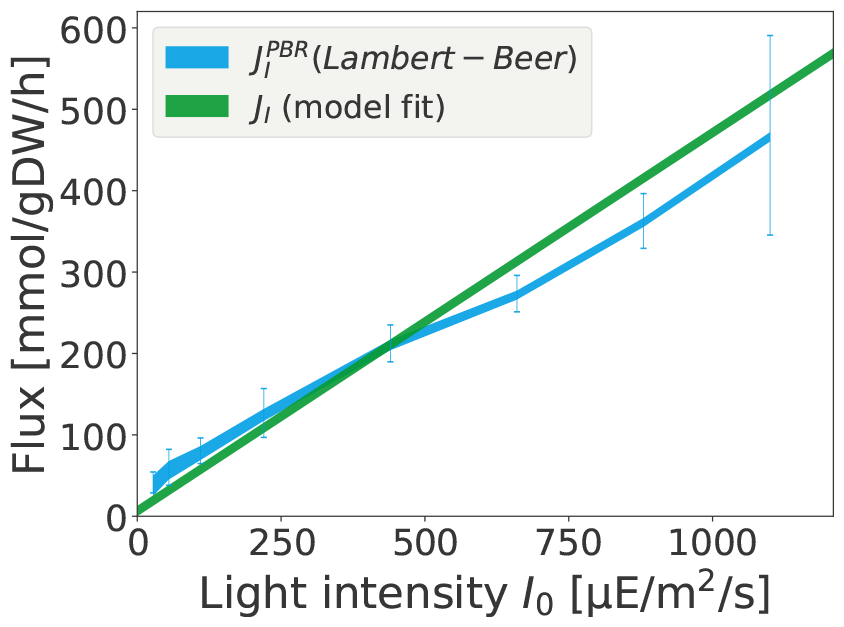
Evaluating light absorption with a mechanistic model of light uptake. Shown is the fitted photon uptake rate *J*_*I*_, estimated according to equation (7) in comparison with the value 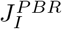 obtained by equation (8). The contribution from blue photons is shown as a (small) shaded area. The value *J*_*I*_ is fitted using only the measured growth-irradiance curve. The value 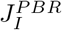 accounts for reactor geometry and its estimation requires knowledge of transmitted light and areal biomass density. Both values are in good agreement. Error bars in 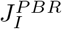 primarily reflect uncertainty in the estimation of the areal biomass density.

### 2.8 Estimating the biomass yield

The estimated rate of photon absorption allows us to assess the biomass yield per photon, defined as the growth rate divided by the photon absorption rate (units: gCDM/mol_*photons*_). Figure 5 shows the biomass yield as a function of light intensity, and compares the values obtained for the experimental growth rates to the values estimated from the model and to the maximal stoichiometric yield obtained by conventional FBA. The latter does not take photoinhibition and light saturation into account, but still accounts for the additional physiological constraints given in Table 1 (hence the biomss yield is slightly lower the maximal yield obtained from the stoichiometric overall equation reported above).

**Fig. 5:**
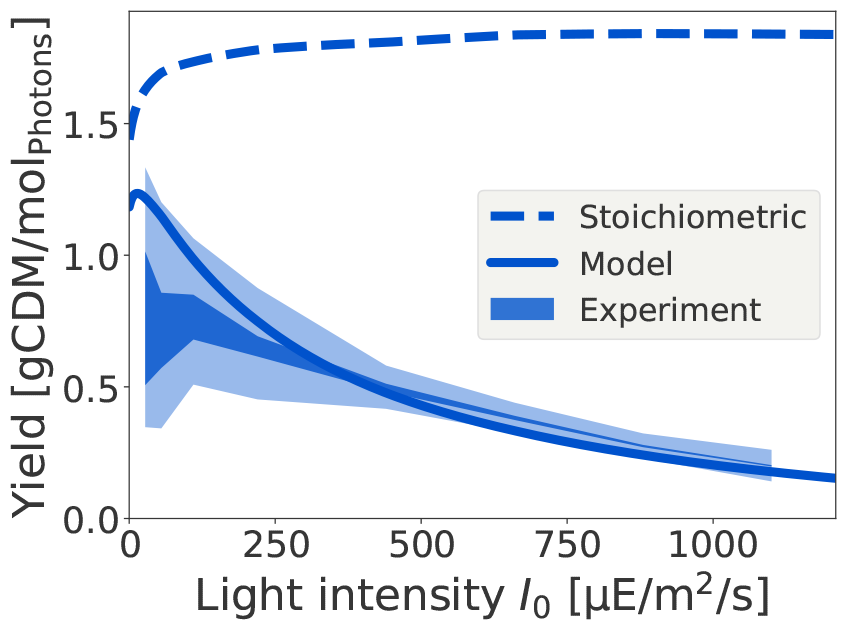
The biomass yield as a function of the incident light intensity. Shown is the stoichiometric biomass yield, as obtained by conventional FBA. Due to non-growth associated ATP utilization, the value of the stoichiometric yield decreases for low light intensities but saturates for higher light intensities. In contrast, the yield obtained from the fitted model (blue line) decreases with increasing light intensity. The experimental yield is depicted as a shaded blue area that reflects the weight of blue photons from 0 to 100%. Error bars are depicted as a light blue area.

The experimental yield (shown as dark blue area) is defined as the measured growth rate divided by the rate 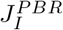 of light absorption, the contribution from blue light is again shown as a shaded area (dark blue), the error intervals are indicated in light color. The biomass yield obtained from the model (solid line) corresponds to the fitted growth rate divided by the light uptake estimated according to equation (7). The simulation takes all physiological constraints, including ATP maintenance, into account. The drop in yield for low light intensities is due to the non-growth associated maintenance requirements.

The difference between experimental and model-derived yield is primarily due to differences in the amount of light absorption at low light intensities (see also Figure 4). The estimated values are significantly below the maximal stoichiometric biomass yield, and are in good agreement with values previously reported for *Synechocystis* sp. PCC 6803. For example, Touloupakis et al. [46] report a biomass yield of *Y*_BM_ ≈ 1.0 gCDM / mol (photons) for *Synechocystis* sp. PCC 6803 in continuous cultures using a light intensity of *I*_0_ ≈ 150 *µ*mol (photons)/m^2^/s.

It is noted that a decreasing biomass yield for higher light intensities is an important property to correctly estimate the expected phototrophic productivities in biotechnological applications [31].

### 2.9 Predicting physiological properties: O_2_ evolution

Using the estimated parameters, the model allows us to evaluate further physiological properties and exchange fluxes. Of particular interest is the oxygen evolution rate in dependence of the light intensity, a key property of oxygenic photosynthesis. Net O_2_ evolution, as reported in Zavřel et al. [1], consists of the contribution from PSII (gross O_2_ production by PSII) minus the use of O_2_ for respiration, Mehler-like reactions, and uptake and release of molecular O_2_ as a stoichiometric substrate in metabolism. We note that, as yet, the data on light-dependent net O_2_ evolution (Figure 2D) was not used in the estimation of growth parameters, and hence can serve as a test of the predictive value of the model.

Figure 6 shows the experimentally measured net O_2_ evolution in comparison with the predicted net O_2_ evolution from the parameterized model. Simulations were performed with the light-dependent BOF. We emphasize that there is no variability in model predictions, that is, given the fitted parameters *α, α*_*b*_, *K*_*L*_, and *k*_*d*_, the rate of O_2_ evolution is fully determined. We further performed a feasibility analysis whether parameters exist that would allow us to exactly match the measured growth rate while constraining the O_2_ evolution to measured values. Such parameters are not feasible.

**Fig. 6:**
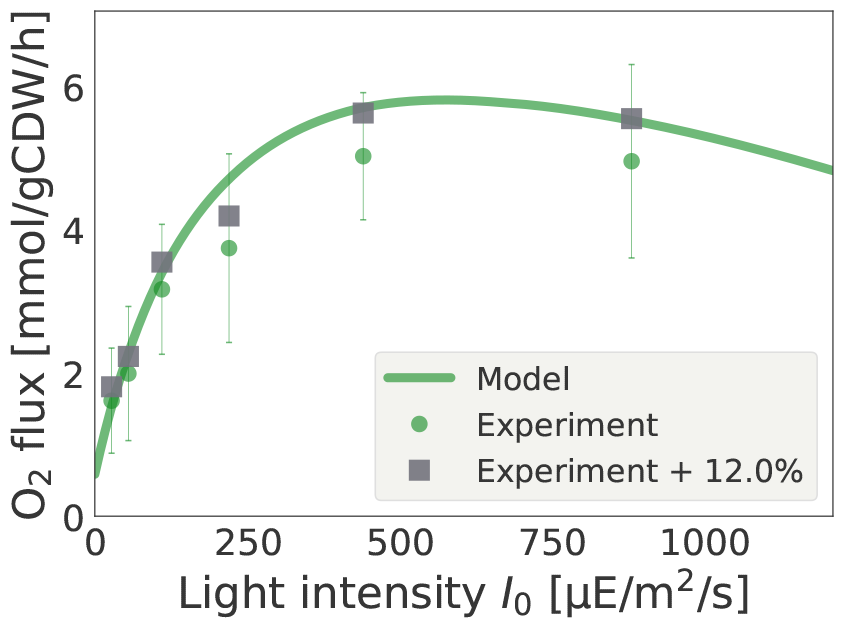
Net oxygen (O_2_) evolution as a function of the incident light intensity. Experimental values (green dots) are compared to the predictions from the fitted model (solid green line). The latter slightly overestimates net O_2_ production. A possible reason for the deviation is a loss of O_2_ via the headspace. The grey squares indicate an offset of ≈ 12% (the values are not fitted and only serve as a guide for the eye).

As shown in Figure 6, the model slightly overestimates O_2_ production in comparison to the experimental data. The difference, while still within estimated error intervals, may stem from several factors, including potential inaccuracies in the metabolic reconstruction, as well as potential systematic errors in the experimental procedures. In particular, following the protocol described in [1, 43], experimental O_2_ evolution was determined by stopping continuous gas supply (bubbling) and subsequently measuring accumulation of dissolved oxygen (dO_2_) in the photobioreactor cuvette. The protocol neglects loss of O_2_ via the headspace during the time of the measurement, and hence slightly underestimated actual O_2_ evolution. As shown previously for the volatile product ethylene synthesized by cells containing a heterologous ethylene-forming enzyme, loss of product via the headspace can shift the measurement by up to 12% [47]. The headspace-liquid partition coefficients (*K*, the ratio of the concentration of molecules between the two phases when at equilibrium) of ethylene and oxygen are similar (*K* = 0.036 and *K* = 0.03, respectively [48, 49]), hence we can expect a similar effect for oxygen. Figure 6 (gray squares) illustrates the effects of a 12% systematic shift in O_2_ evolution that would explain the observed difference. We note that, in addition to O_2_ loss via headspace, also other factors may play a role.

We further emphasize that using the light-dependent adjustment of the biomass composition (BOF) is crucial for the prediction of O_2_ production. Supplementary Figure S2 shows the predictions for the static BOF used in previous reconstructions [5, 24]. The results support the recent analysis of Dinh et al. [40] that, while the prediction of the growth rate itself does not depend crucially on the definition of the BOF, individual reaction fluxes can be highly dependent.

### 2.10 Electron transport and the ATP/NADPH ratio

In addition to O_2_ exchange, the fitted model can be used to investigate the interplay between cyclic (CET) and linear (LET) electron transport and, more generally, the electron flow of PSII compared to PSI and the resulting ratio of synthesis of ATP relative to NADPH. Maximizing synthesis of biomass with the given constraints gives rise to a required (optimal) ratio of generated ATP relative to NADPH. This optimal ratio exceeds the ratio synthesized by LET and requires a contribution from CET to meet additional ATP demands.

We note that calculating the required (optimal) ATP/NADPH ratio in the model exhibits variability since the redox carrier NADPH can, to some extent, be substituted by other carriers, such as reduced ferredoxin [50]. The use of alternative redox carriers is stoichiometrically equivalent and cannot be distinguished by stoichiometric constraint-based analysis. We therefore employ flux variability analysis (FVA) to evaluate the (range of) flux values of the ATPase and the ferredoxin-NADP(+) reductase (FNR) reaction. FVA allows us to estimate the predicted range of (optimal) ATP/NADPH synthesis rates as a function of light intensity in the fitted model. The results are shown in Figure 7, with the variability indicated as a shaded area. The lower bound corresponds to the use of NADPH as primary redox carrier, i.e. using NADPH whenever stoichiometrically possible.

**Fig. 7:**
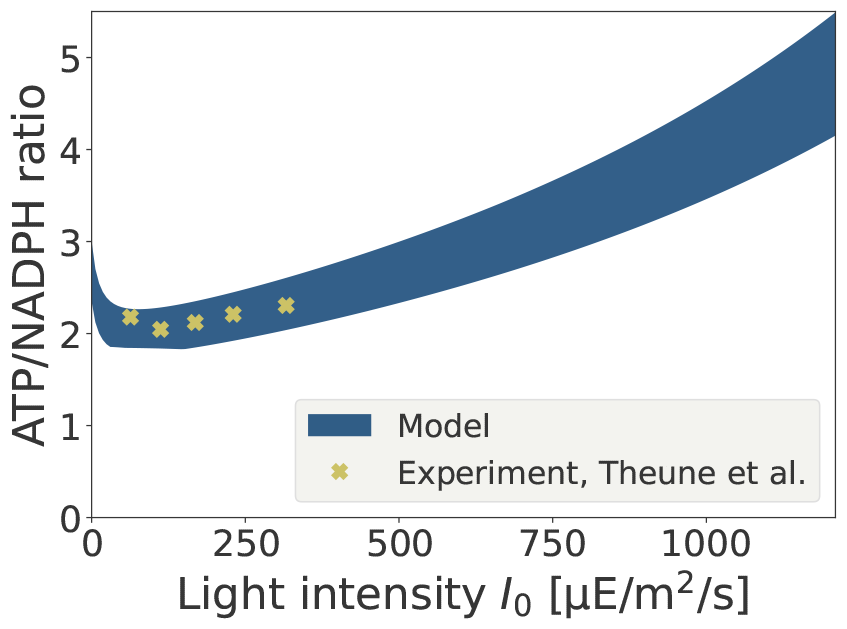
Predicted range of (optimal) ATP versus NADPH synthesis estimated from the fitted model, shown as shaded area. The variability is due to substitution of NADPH by other redox carriers, the lower bound corresponds to the case with NADPH as the primary redox carrier. The predicted values are in good agreement with experimental values reported by Theune et al. [32].

The predicted values can be compared to recent experimental estimates reported by Theune et al. [32], who quantified CET and the resulting ATP/NADPH ratio in *Synechocystis* sp. PCC 6803. The experimental values are shown as dots in Figure 7. We note that the experimental setup of Theune et al. [32] differs from the setup used in Zavřel et al. [1], hence the reported light intensities are not necessarily directly comparable (the impact of light intensity also depends on the volumetric biomass density and the reactor geometry). Nonetheless, the predicted and experimental values are in good agreement, indicating that the fitted model provides a reasonable description of the energetics of cyanobacterial growth under these conditions. As a test for further analysis, the model suggests that the ATP/NADPH ratio will increase with increasing light intensities, primarily due to the increased requirement of repair mechanisms to alleviate photodamage. A comparison of the electron fluxes (CET versus LET) compared to the data from Theune et al. [32] is provided as Supplemental Figure S3.

## 3 Conclusions

In this work, we presented a quantitative analysis of light-limited cyanobacterial growth based on an updated genome-scale reconstruction of the model strain *Synechocystis* sp. PCC 6803. The revised reconstruction is provided as an SBML file. In addition to the full model, we also provide a stoichiometrically reduced core model (Supplementary File S4) using an established algorithm for network reduction [28]. The reduced core network is useful for applications that benefit from a smaller reaction network that nonetheless allows to simulate cyanobacterial growth with a stoichiometrically correct mass balance.

While constraint-based analysis of cyanobacterial metabolism is well established, in particular in the context of strain design and biotechnological applications [24, 51, 52], a quantitative analyses of cyanobacterial growth using genome-scale models are still scarce. In particular, compared to a stoichiometric description of heterotrophic growth, the description of phototrophic metabolism and growth gives rise to additional challenges due to the utilization of light as a primary source of energy. As yet, most constraint-based analyses of phototrophic metabolism consider photons analogous to nutrient molecules, an approach that fails to capture key properties of photosynthesis, such as a variable quantum yield or photodamage.

To address this challenge, our work proposes a novel method to describe light absorption and light utilization in constraint-based models of phototrophic growth. Our approach is similar to methods that consider saturable nutrient uptake rates for heterotrophic organisms [53], but is specifically tailored to describe photosynthetic light absorption. Our method is motivated by mechanistic models of photosynthesis, and the required parameters have a clear interpretation in the context of phototrophic growth. In the simplest case, our approach makes use of 3 additional parameters, a light absorption coefficient *α*, a maximal photosynthetic capacity *K*_*L*_, and a rate constant *k*_*d*_ that describes photodamage. As shown in this work, these parameters reflect the three parameters of the phenomenological Haldane/Aiba equation and can be estimated from a experimentally obtained growth-irradiance curve (Figure 2A). Within such an estimation, the value of the incident light intensity can be considered as a descriptive property of the culture condition, detailed knowledge about light quality or reactor geometry are not required. Constrained-based optimization is then based on (an upper limit on) the rate 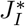 of productively used photons.

As demonstrated, the method proposed here gives rise to a nonlinear dependence of the growth rate on the incident light intensity (Figure 3). The parameterized model allows for analyses that go beyond conventional FBA, and includes a (decreasing) photosynthetic efficiency and (decreasing) biomass yield for increasing light intensity (Figure 5). The latter properties are relevant in the context of biotechnological applications and reactor design [31, 39, 54], and are difficult to address using conventional FBA. Supporting previous works [40], our analysis also indicates that the cellular growth rate does not crucially depend on the detailed definition of the BOF, whereas prediction of individual fluxes can be sensitive to the assumed intracellular composition.

Furthermore, the model is capable to quantitatively predict several physiological properties, such as the net O_2_ evolution (Figure 6) and the required (optimal) ATP/NADPH ratio (Figure 7), without additional assumptions or additional fitting of parameters. In particular, the ratio of ATP to NADPH synthesis is a key property for biotechnological applications. Most heterologous products have an ATP/NADPH requirement that is lower than the ratio required for the synthesis of cellular biomass [24, 51].

We emphasize that, as shown in Figure 7, our approach may give rise to changes in intracellular properties, such as a variable ATP/NADPH ratio, as a function of light intensity–such changes can not arise within the conventional strictly linear framework of FBA. Nonetheless, our approach retains most advantages of conventional FBA in terms of computational simplicity. In particular, while it is possible to devise more complex methodologies to describe light absorption and phototrophic growth, e.g., genome-scale ME/RBA models [11] or hybrid models that combine a detailed kinetic description of photosynthesis with a stoichiometric model of metabolism [19], such models also require a significant additional investment with respect to the required knowledge of parameters, and give rise to a significantly increased computational burden. In contrast, the strength of our approach is that it keeps the computational and conceptual simplicity of conventional FBA, while still allowing for a quantitative description of growth properties. We conjecture that, despite the expected increasing availability of RBA/ME-type models also for cyanobacterial strains [11], FBA will remain a method of choice for constraint-based analyses in the foreseeable future.

While our focus was a quantitative description of light-limited phototrophic growth based on the experimental setup described by Zavřel et al. [1], our approach can be generalized and applied also in different contexts. Applications not considered here include, for example, limitation by other environmental factor than light intensity and the exudation of organic compounds [29]. A further important development would be a generalization to arbitrary light spectra and their impact on growth [7, 42, 52, 55]. Suitable ways to include different light qualities would be a wavelength-dependent absorption, as employed here with respect to the blue background illumination, as well as separate saturation functions for photosystem II and photosystem I. A detailed parameterization of such a generalization, however, is currently not possible due to insufficient availability of quantitative data.

We argue that the approach presented here advances the quantitative analysis of light-limited phototrophic growth, and will be readily applicable in many applications that currently make use of conventional FBA. In particular, our approach demonstrates that key properties of photosynthesis, such as a variable quantum yield and photodamage, can be incorporated into established constraint-based models of phototrophic growth without sacrificing computational simplicity.

## 4 Materials and Methods

### 4.1 The metabolic network of *Synechocystis* 6803

The metabolic reconstruction of *Synechocystis* sp. PCC 6803 is based on previous reconstruction [5, 24], and revised according to current literature and other recent reconstructions [8]. SBO annotation was added using the SBOannotator version 0.9 [56]. Compared to [24], added reactions relate to the Entner–Doudoroff pathway, tyrosine and phenylalanine biosynthesis via prepheneate, biotin metabolism, methanol detoxification, and purine and pyrimidine metabolism. A description of all changes can be found in Supplementary File S5. We note that recent additions to reconstructions of the metabolism of *Synechocystis* sp. PCC 6803 have no major impact on the results of this study. The model is encoded in SBML format (SBML Level 3 Version 1) and is available as Supplementary File S2 (SBML) and Supplementary File S3 (xlsx). A static biomass objective function (BOF) adopted from [5, 24] is used as a reference, the stoichiometric coefficients of macromolecular components are provided in Supplementary Table S1.

### 4.2 Flux balance analysis (FBA)

FBA utilizes linear programming, a mathematical technique for constraint-based optimization [17, 18]. FBA seeks to find a vector *ν* of reaction fluxes that satisfies the constraints (mass balance and flux constraints) while maximizing a (linear) objective function *ν*_*BOF*_. The canonical form of FBA is

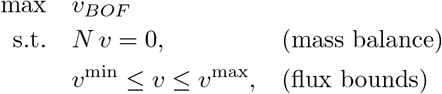

where the biomass objective function *ν*_*BOF*_ = *c*^*T*^ *ν* is a linear combination of reaction fluxes, and *ν*^min^ and *ν*^max^ denote upper and lower bounds on the reaction rates, respectively. In our simulations 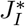 serves as an upper bound for the photon uptake rate, corresponding to the sum of photon utilization rates at photosystem I and II. Uptake of other nutrients is not constrained.

In this work, COBRApy [57], version 0.29.0, with the GLPK (GNU Linear Programming Kit, version 5.0) solver was used. Scripts and further instructions are available as Supplementary File S5.

### 4.3 Stoichiometric network reduction

In addition to the full metabolic network, we derive a reduced core model using established methods of stoichiometric network reduction [28]. Stoichiometric network reduction requires to specify a set of reactions and metabolic functions that are protected. In our case, we focus on key reactions of central metabolism, such as those associated with the Calvin-Benson cycle and Photosynthesis, amounting to a total of 75 reactions. Additionally, we protect the capability of phototrophic growth using atmospheric CO_2_ and light as source of carbon and energy, respectively. Specifically, we require the network to grow with at least 99% of the rate of the full network, given a maximal photon uptake rate of 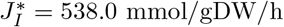,

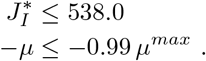

Network reduction then proceeds in two steps: firstly, reactions which are not protected and are not required to sustain the protected metabolic functionality are iteratively removed (pruning). Secondly, the remaining reactions are lumped into overall reactions whenever possible, i.e., without loosing functionality (compression). The algorithm is described in [28].

The script for network reduction was developed in MATLAB, employing NetworkReducer tool from the CellNetAnalyzer package (v2023.1) [58]. The resulting pruned and compressed model has a significantly lower dimension compared to the original GSMR, consisting of 87 internal metabolites and 97 reactions. The reduced network is available as Supplementary File S4.

### 4.4 Light utilization and variable quantum yield

We propose a novel approach to incorporate light absorption and utilization into constraint-based models of light-limited phototrophic growth. To motivate our approach, we recall that within conventional FBA, the nutrient-limited maximal growth rate can be described by the product of the uptake rate *J*_*I*_ and the maximal biomass yield 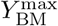 (in the absence of additional constraints),

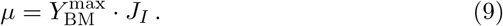

The maximal biomass yield 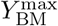 depends on the stoichiometry of the metabolic reaction network.

To incorporate light absorption and a variable quantum yield, we first consider the dependence of the growth rate on the incident light intensity (units mol photons/m^2^/s) using a phenomenological Monod equation,

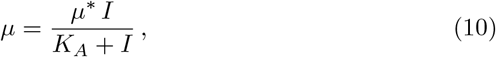

where *µ*^*∗*^ denotes the maximal growth rate, and *K*_*A*_ the half-saturation constant. The Monod equation can be (mathematically equivalent) rewritten as [31]

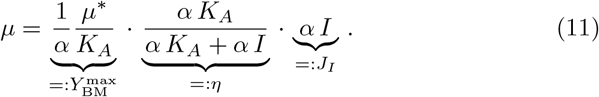

where the term 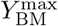 denotes the maximal biomass yield per mol photons (see below). Using the definition *K*_*L*_ := *α K*_*A*_, we obtain

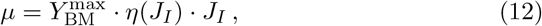

with

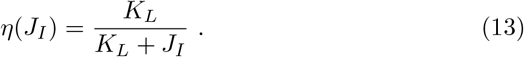

As noted previously [31], the light-limited growth rate of a phototrophic organism is a product of the rate of light uptake *J*_*I*_, the (dimensionless) quantum yield or photosynthetic efficiency *η*, and the maximal biomass yield per photon 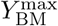.

In analogy to equation (9), we therefore make use of an effective light absorption rate 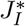 that incorporates the decreasing quantum yield as a function of light absorption,

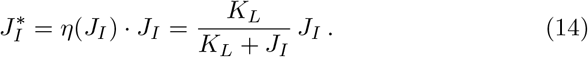

As outlined in the main text, the parameter *K*_*L*_ (units: *µ*E/gCDM/s) corresponds to a maximal capacity of light utilization. Under low light conditions, *J*_*I*_ ≪ *K*_*L*_, almost all absorbed photons are used productively and 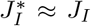, whereas for high light intensities, *J*_*I*_ → ∞, an upper limit 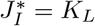 is reached. The latter also justifies the definition of 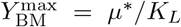 as the maximal biomass yield.

Equation (14) is then utilized to incorporate the variable quantum yield of photosynthesis into constrained-based models of phototrophic growth. Required parameters are *K*_*L*_ and *α*. We note that our approach, similar to previous coarse-grained models [33, 34, 39], considers the quantum efficiency of the complete photosynthetic electron transport chain and does not distinguish between photosystem I and II.

### 4.5 Derivation using a 2-state model of photosynthesis

Our approach can be further motivated by a mechanistic 2-state model of photosynthesis. Similar to kinetic coarse-grained models of photosynthesis [33, 34, 39], we consider the activation of a photosynthetic unit (representative of the entire photosynthetic electron transport chain) by the absorption of a photon *P* ^0^ → *P*^*∗*^ with a rate *ν* _1_ = *σ* ·*P* ^0^ ·*I*. The rate depends on an effective absorption area *σ* per photosynthetic unit (unit: area/mol), the concentration of inactive (or open) photosynthetic units *P* ^0^ (unit: mol/gCDM), and the light intensity I. The relaxation *P*^*∗*^ → *P* ^0^ back to the open state provides energy for the cell and occurs with a rate *ν* _2_ = *k*_2_ *P*^*∗*^. The growth rate can then be described by the product of *ν* _2_ with the biomass yield per unit energy (photon), 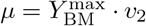

Given these rate equations, the steady-state concentration of activated *P*^*∗*^ is

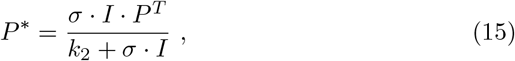

where *P* ^*T*^ = *P* ^0^ + *P*^*∗*^ denotes the total concentration of photosynthetic units. Hence, using the definitions *K*_*L*_ = *k*_2_ · *P* ^*T*^ (corresponding to the maximal capacity of the photosynthetic units) and *J*_*I*_ = *σ* · *I* · *P* ^*T*^ (corresponding to the total rate of light absorption), the specific growth rate is

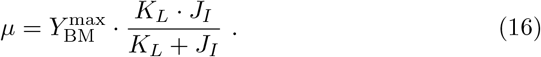

Again we obtain an effective rate 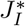 of light absorption,

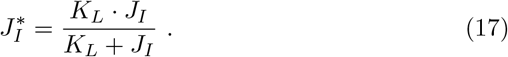

We note that in this view, the maximal capacity is given by the total expression of the photosynthetic electron transport chain multiplied with its rate constant (lumped into a single reaction), likewise light absorption depends on the expression of the photosynthetic apparatus with *α* = *σ P* ^*T*^.

### 4.6 Parameter estimation

Estimation of parameters was performed using the nonlinear least-square algorithm from SciPy (v. 1.11.3) <monospace>Optimize</monospace> [59]. Handling of SBML files and FBA simulations are carried out with COBRApy (v. 0.29.0) [57].

In brief, the unknown parameters are *α, α*_*b*_, *K*_*L*_, and *k*_*d*_, as well as 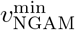 and 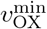. For each light intensity, the parameters *α, α*_*b*_, *K*_*L*_ determine the effective light uptake rate 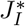. The values of 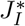, as well as the parameters *k*_*d*_, 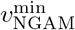 and 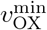 are then part of the linear program to maximize the BOF.

The nonlinear least-square algorithm seeks to minimize the difference between the maximal growth rate obtained from the model and the experimental growth across all light intensities, as well as to minimize the difference between the rate of non-light associated O_2_ uptake in the model compared to the experimentally determined values (Figure 2A and E). Within the least-square algorithm, the differences in growth rate are weighted with a factor 10^3^. For each light intensity, the biomass objective function is adjusted to reflect the experimentally measured values of glycogen and protein macrocomponents at the particular light intensity. The rate of light-dependent O_2_ (Figure 2D) is not used in the fitting process. Python scripts are provided as Supplementary File S6.

### 4.7 Model evaluation: O_2_ evolution and ATP/NADPH ratio

Given the fitted parameters, the model is evaluated for the full range of light intensities. Simulations are conducted for red light intensities ranging from 0 to 1210 *µ*E/m^2^/s, using evenly spaced data points (in total 1000 data points). Maximizing the BOF for each light intensty results in estimations for metabolic fluxes and O_2_ export, as well as O_2_ production at PSII.

To determine the ratio of ATP/NADPH synthesis, the fluxes of the thylakoid membrane complexes ATPase and ferredoxin-NADP(+) reductase (FNR) were evaluated. Due to the variability of the FNR reaction, flux variability analysis (FVA) was employed to obtain the minimal and maximal values of these fluxes for each (red) light intensities ranging from 0 to 1210 *µ*E/m^2^/s (200 data points). The ratio of ATP/NADPH synthesis was then calculated as 3*J*_ATPase_*/J*_FNR_, due to the stoichiometry of the ATPase with 3 ATP per full cycle.

### 4.8 Sensitivity analysis

The impact of the estimated parameters on the growth rate is evaluated using sensitivity analysis [17]. For any parameter *k* we calculate the relative difference in the maximal growth rate given a small change in one of the estimated parameters. Formally,

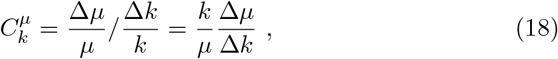

where *k* stands for the value of any of the estimated parameters. The scaled or normalized sensitivities 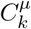 specify the relative change in the (maximal) growth rate upon a relative change in the parameter *k*. Positive values imply the growth rate increases with an increasing value of the parameter.

Computationally, the dimensionless (scaled or normalized) relative sensitivities 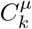 were evaluated symmetrically around each value using a 2% change in the parameter, that is, Δ*k* = *k*^+^− *k*^*−*^ with *k*^±^= *k* ± 0.01 · *k*. Subsequently the resulting change in maximal growth rate was evaluated. Results of the sensitivity analysis for the parameters *α, α*_*b*_, *K*_*L*_, and *k*_*d*_, as well as for the light intensity *I*_0_ are shown in Figure 8.

**Fig. 8:**
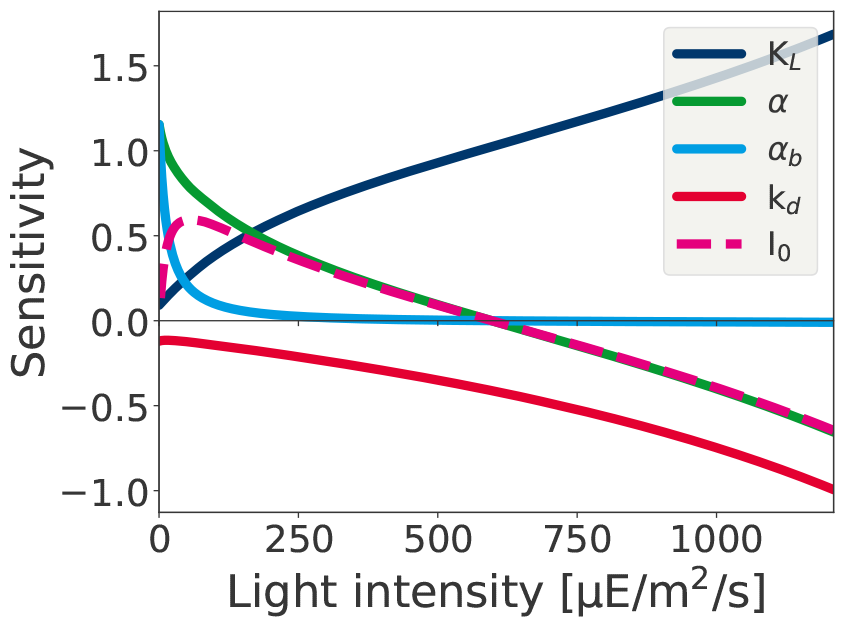
Normalized sensitivity analysis of the growth rate with respect to (small) changes in estimated parameters *α, α*_*b*_, *K*_*L*_, and *k*_*d*_. The plot illustrates the different growth regimes. For low light intensities, the coefficients describing light absorption have the largest impact (light-limited growth). The sensitivity with respect to the constant blue background illumination (*α*_*b*_) rapidly drops. For high light intensities, the impact of the light absorption coefficients and the light intensity *I*_0_ becomes negative. In the regime of photoinhibition, the impact of *k*_*d*_ and the impact of the capacity *K*_*L*_ is high.

The changes in scaled sensitivities 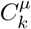 reflect the different growth regimes. For low light intensities, the light absorption parameters *α* and *α*_*b*_, as well as the light intensity *I* itself, have the highest impact on growth. The impact of the parameter *α*_*b*_ rapidly drops due to the low constant contribution of blue light. With increasing light intensity, the impact of the maximal capacity *K*_*L*_ and the impact of the rate constant of photodamage *k*_*d*_ increases (the impact of *k*_*d*_ is always negative, i.e., increasing *k*_*d*_ will always lower the maximal growth rate). At the optimal light intensity 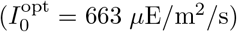 the sensitivity with respect to the absorption parameter and the light intensity is zero (at this point, the contribution from blue light has no discernible impact anymore). For even higher light intensities, the cell is in the regime of photoinhibition, i.e., the impact of light intensity on the growth rate is negative and the parameter *k*_*d*_ has a high impact.

## Supplementary Text and Figures

**Table S1:** A static biomass objective function (BOF) adopted from [5, 24]. The static BOF is used as a reference. Shown are the amounts of coarse-grained components.

| Stoichiometry | Metabolite | Name |
| --- | --- | --- |
| -0.51 | bm_pro_c | Protein component of biomass |
| -0.02 | bm_pigm_c | Pigment component of biomass |
| -0.03 | bm_sol_c | Soluble pool component of biomass |
| -0.01 | bm_ion_c | Ion component of biomass |
| -0.17 | bm_rna_c | RNA component of biomass |
| -0.12 | bm_memlip_c | Lipid component of biomass |
| -0.06 | bm_cw_c | Cell wall component of biomass |
| -53.35 | atp_c | ATP |
| -53.35 | h2o_c | H<sub>2</sub>O |
| -0.03 | bm_dna_c | DNA component of biomass |
| -0.03 | bm_glycogen_c | Glycogen component of biomass |
| 1.0 | biomass_e | Biomass |
| 53.35 | adp_c | ADP |
| 53.35 | pi_c | Orthophosphate |

**Fig. S1:**
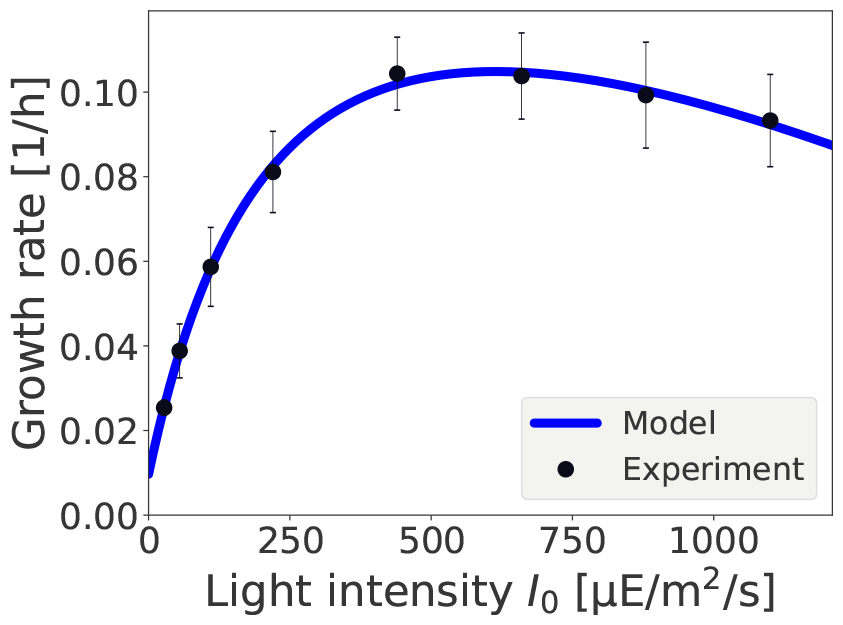
A fit of the growth rate using the static BOF. Methods identical to as Figure 3 but with a static BOF for all light intensities. The detailed composition of the BOF has no major impact on the quality of the fit of the model to the measured growth rate.

**Fig. S2:**
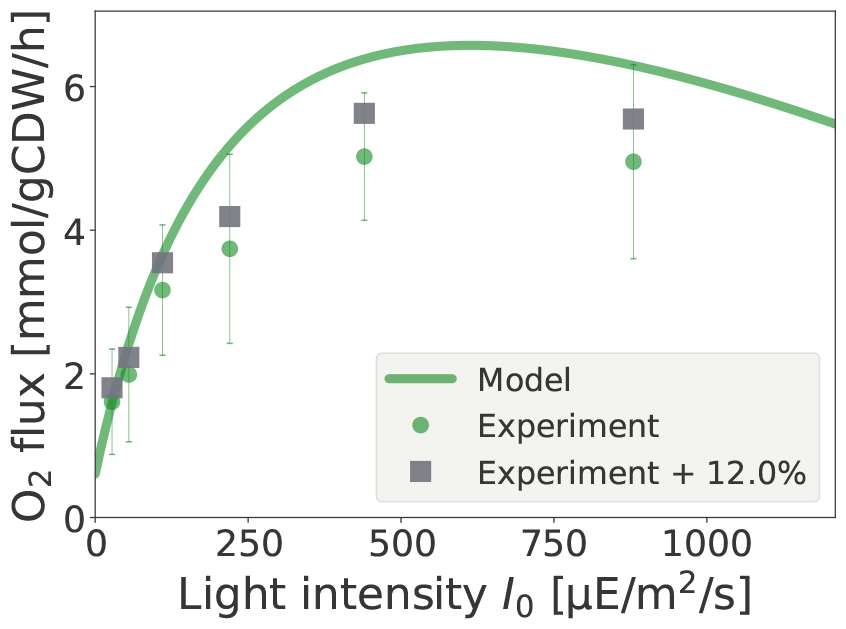
Prediction of O_2_ evolution obtained from the model compared to experimental results using the static BOF. Compare with Figure 6. The detailed composition of the BOF has impact on the quality of the prediction of O_2_ evolution. The grey squares indicate a deviation of 12% from the predicted values.

**Table S2:**
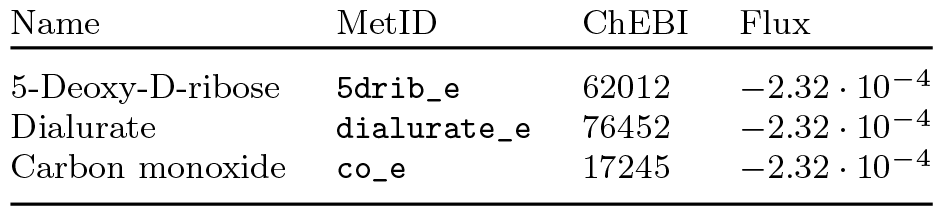
According to the model, growth results in the synthesis and excretion of three inevitable byproducts. The table provides the products and their excretion flux rate (in units mmol/gCDM/h) for a reference growth rate of 1 h^*−*1^ and a reference CO_2_ fixation flux of 41.4 mmol/gCDM/h.

**Fig. S3:**
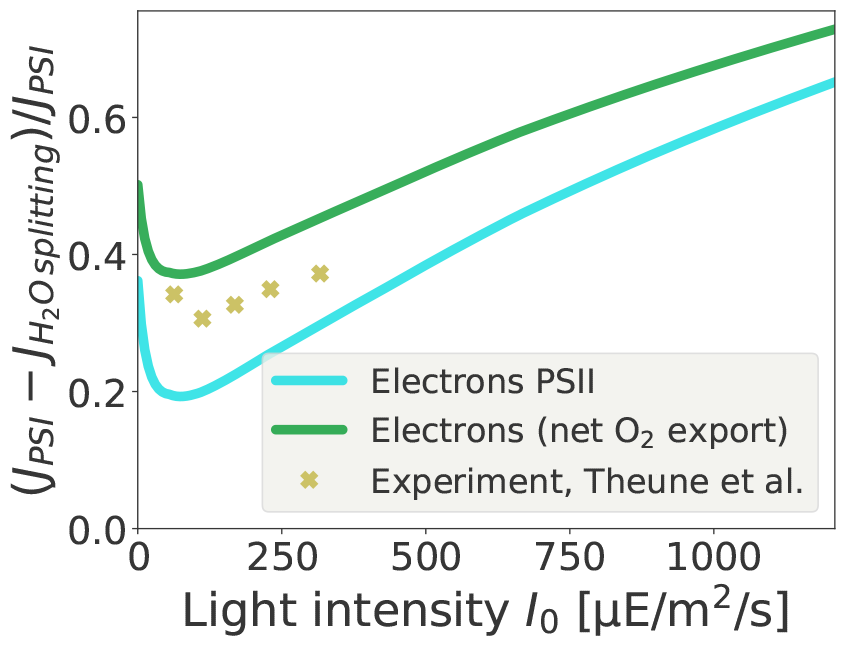
Predicted linear (LET) versus cyclic electron transport (CET) as described in the main text. Electrons from PSII (water splitting) are counted either directly from the flux at PSII (lower line) or are estimated from net *O*_2_ export (4 electrons per *O*_2_, upper line). The latter (upper line) corresponds to the experimental quantification of Theune et al. [32]. We note that light intensities in both setups cannot necessarily be directly compared, since the effective light intensity also depends on culture density and vessel geometry, hence the light axis may be shifted.

**Fig. S4:**
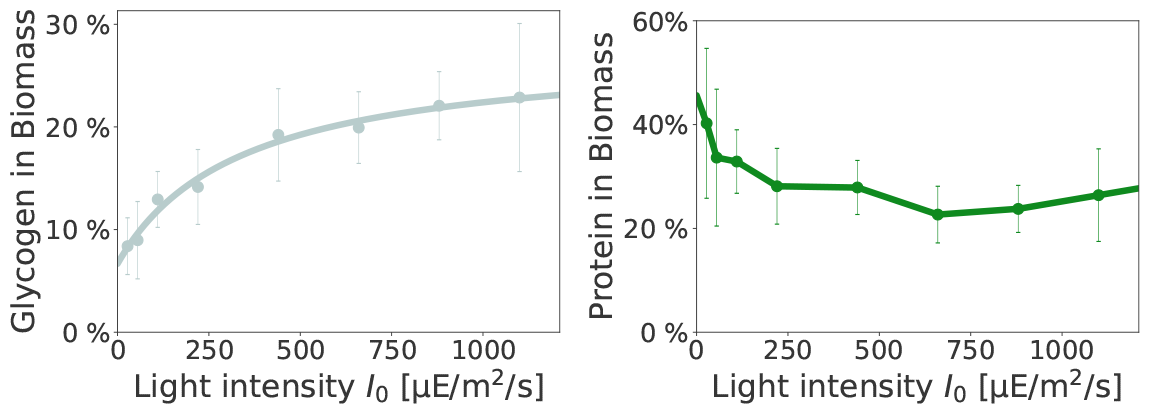
The experimentally determined glycogen and protein mass fractions in the BOF. For a model evaluation across the entire range of light intensities, the glycogen mass fraction was fitted to a Monod function with offset, i.e., *glyc* = *a I*_0_*/*(*b* + *I*_0_) + *c* with *a* = 0.21, *b* = 336.56, *c* = 0.07. The protein mass fraction was linearly interpolated between experimental values. All data as originally reported in [1].

**Fig. S5:**
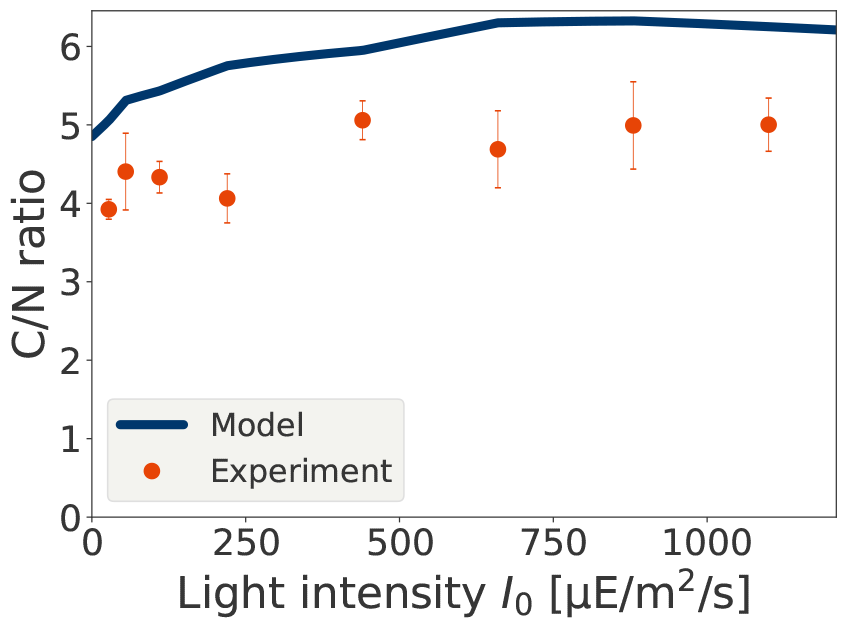
Comparison of the C:N ratio obtained from the model versus experimental values reported by Zavřel et al. [1].

## Notes

### Competing Interest Statement

The authors have declared no competing interest.

